# SYNY: a pipeline to investigate and visualize collinearity between genomes

**DOI:** 10.1101/2024.05.09.593317

**Authors:** Alexander Thomas Julian, Jean-François Pombert

## Abstract

Investigating collinearity between chromosomes is often used in comparative genomics to help identify gene orthologs, pinpoint genes that might have been overlooked as part of annotation processes and/or perform various evolutionary inferences. Collinear segments, also known as syntenic blocks, can be inferred from sequence alignments and/or from the identification of genes arrayed in the same order and relative orientations between investigated genomes. To help perform these analyses and assess their outcomes, we built a simple pipeline called SYNY (for synteny) that implements the two distinct approaches and produces different visualizations. The SYNY pipeline was built with ease of use in mind and runs on modest hardware. The pipeline is written in Perl and Python and is available on GitHub (https://github.com/PombertLab/SYNY) under the permissive MIT license.

## Introduction

The identification of collinear segments (also known as syntenic blocks) between genomes is a common task in comparative genomics. Albeit traditionally used in genetics to refer to genes that are located on the same chromosomes but not necessarily in the same arrangement, the term synteny —from the Greek words *syn* and *ταινια* meaning *on the same band*—has been repurposed in genomics to refer to genes that are arrayed in the same order and relative orientations between genomes [1]. The positional information retrieved from collinearity inferences can be used for several purposes. For example, it can be used to help detect genes that might have been overlooked during genome annotation, it can be used to help identify which genes are real orthologs within a pool of similar paralogs retrieved from sequence homology searches, and it can be used to develop and test various evolutionary hypotheses such as reconstructing the structure and content of ancestral genomes [2,3].

In comparative genomics, collinearity assessments are usually performed by sequence alignments and/or by looking for sets of genes, *i*.*e*. clusters, that share the same order and relative orientations between genomes. Since alignment-based approaches do not rely on genome annotations, they can be performed on unannotated sequences, but because these approaches rely on the presence of a modicum of homology between the sequences being compared, they tend to struggle when confronted with high levels of divergence. Conversely, while approaches based on gene clusters require annotated sequences (and thus struggle with poorly annotated genomes), they tend to fare better with high levels of divergence. Hypervariable intergenic regions are not considered in gene cluster analyses and by further restricting the analyses to protein-coding genes, searches can be performed across larger evolutionary distances by leveraging the amino acid sequences of their products. Amino acid sequences are not affected by silent mutations nor by codon usage biases, they feature a larger-state space than nucleotide sequences, and their back mutation probabilities are lower than those from nucleotides [4].

Although several tools have been developed over the years to help infer synteny between genomes (see Liu et al. [5] for a review), no single tool met our comparative genomics needs. Namely, we needed a tool that could produce high quality circular and linear collinearity maps from both genome alignment and gene cluster inferences starting from NCBI GenBank Flat files. To rectify this and allow other research groups to perform similar analyses, we developed SYNY, a simple pipeline to investigate and visualize collinearity between genomes.

### Pipeline overview

The SYNY pipeline was developed on Linux. Its dependencies can be installed automatically on most Linux distributions using *setup_syny*.*pl*. Collinearity inferences with SYNY involve three steps: (i) data parsing; (ii) collinearity inferences from genome alignments and/or gene clusters; and (iii) plotting of the results (Fig. 1). All three steps can be performed using the *run_syny*.*pl* master script. Briefly, this script extracts genome/protein sequences and annotation data from NCBI GenBank (.gbff) input files. It identifies collinear sequences by performing round-robin pairwise genome alignments with minimap2 [6]. It locates collinear protein-coding gene clusters by identifying protein orthologs using bidirectional homology searches with DIAMOND [7], finding gene pairs arrayed identically between genomes (with and/or without gaps), and reconstructing clusters from overlapping gene pairs. It then plots the collinear segments inferred from these approaches as dotplots, chromosome maps (hereby referred to as barplots) and Circos [8] plots. It also summarises the corresponding metrics as heatmaps. All plots are generated in Portable Network Graphic (PNG) and Scalable Vector Graphics (Scalable Vector Graphics) formats.

**Figure 1.**
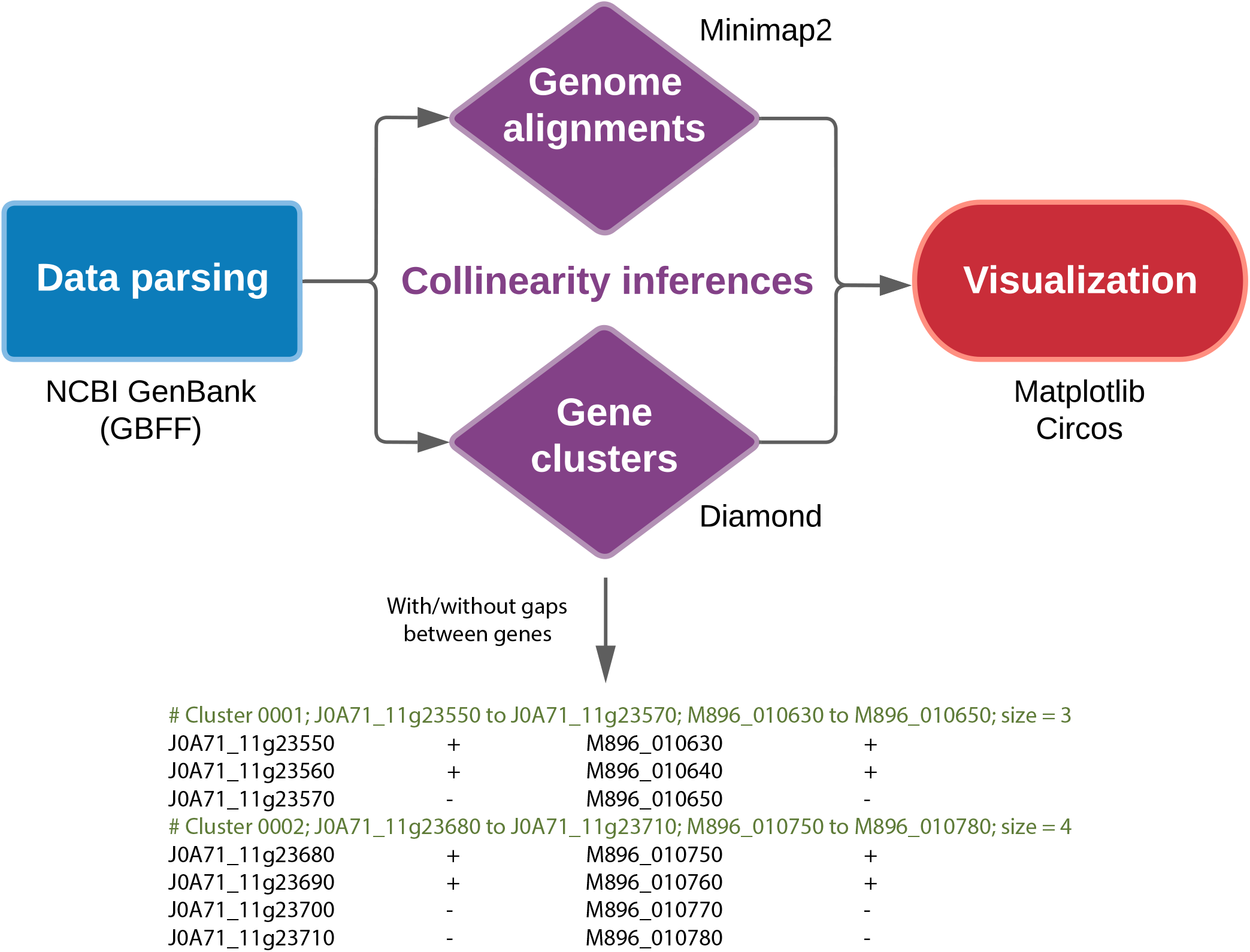
Fig. 1. Overview of the SYNY pipeline. The pipeline contains three main steps: data parsing, collinear inferences, and data visualization. SYNY uses NCBI GenBank Flat files (.gbff) as input. Collinear inferences can be performed from genome alignments and/or from gene clusters. Collinear visualizations include chromosome maps (barplots), dotplots, and Circos plots. Collinear similarities are also summarized as heatmaps.

### Hardware and software requirements

The SYNY pipeline was tested on Linux operating systems (Fedora 40, Ubuntu 22.04.4, and openSUSE Tumbleweed) running as virtual machines (VMs) inside the Microsoft Windows Subsystem for Linux. All tests were performed on a laptop equipped with an Intel Core i5-12500H processor and 12 Gb of RAM allocated to the VMs. The SYNY pipeline is built in Perl and Python. It requires the PerlIO::gzip Perl module and the matplotlib, seaborn, pandas and scipy Python libraries. Minimap2 [6] and DIAMOND [7] are required to perform pairwise genome alignments and sequence homology searches, respectively. Circos [8] is required to generate the corresponding plots.

### Case example – Comparing microsporidia genomes

Microsporidia from the genus *Encephalitozoon* are human-infecting pathogens causing chronic diarrhea, bronchitis, conjunctivitis and/or encephalitis in afflicted patients [9]. With tiny yet complete nuclear genomes totaling less than 3 Mbp, these obligate intracellular pathogens constitute paradigms of genome streamlining in eukaryotes [10]. Their closest known relatives from the genus *Ordospora*, whose members infect crustaceans but not humans, also arbor diminutive genomes [11]. The *Encephalitozoon* and *Ordospora* genomes constitute a good example dataset for the following reasons. Their small sizes make for quick computations, they are highly collinear, and they exhibit a high level of sequence divergence, with average nucleotide identity (ANI) values within the *Encephalitozoon* and *Ordospora* genera of 74-81% and 83%, respectively [11], and between the two genera of about 68% (computed here with OrthoANIu v1.2 [12]). Altogether, the divergence thresholds within (> 15%) and between (> 30%) these genera allow us to demonstrate the limitations of alignment-based collinearity inferences.

Using SYNY with default settings on a total of five genomes (*E. intestinalis, E. hellem, E. cuniculi*, O. *colligata, O. pajunii*) downloaded from NCBI (with Encephalitozoonidae.sh), we were able to infer and plot the collinearity between each genome in less than five minutes (264 seconds) on a laptop equipped with a mobile Intel i5-12500H central processing unit (CPU). This produced a total of 40 dotplots, 40 barplots, and 40 Circos plots: *i*.*e*. pairwise comparisons between queries and subjects are performed in both directions while self comparisons are skipped (5 genomes x [5 genomes – 1] * 2 directions = 40). When allowing three different gap thresholds (0, 1 and 5) for gene clusters inferences and requesting pairwise and concatenated Circos plots using both the normal and inverted karyotypes, the process ran for 13 minutes on the same laptop (from which 11 minutes were for Circos plotting), and produced a total of 80 dotplots, 80 barplots, and 168 Circos plots ([80 pairwise + 4 concatenated plots] * 2 karyotypes).

Examples of collinear plots generated by SYNY from gene clusters and genome alignments are shown in Fig. 2. As expected from the high level of sequence divergence inherent to this dataset, collinear segments inferred from genome alignments (Fig. 2; right) were more fragmented than those reconstructed from gene clusters (Fig. 2; left) in comparisons between species belonging to distinct genera. This artefact was caused by limitations of the genome alignment process, which resulted in as little as 19.1% of the bases mapped in collinear segments between *Encephalitozoon* and *Ordospora* species. In contrast, no less than 66.7% of the protein-coding genes between these species were found in clusters, resulting in much more contiguous collinear plots that more accurately represent the structural similarities between these genomes. When comparing genomes from within the same genera, both approaches produced similar results (not shown).

**Figure 2.**
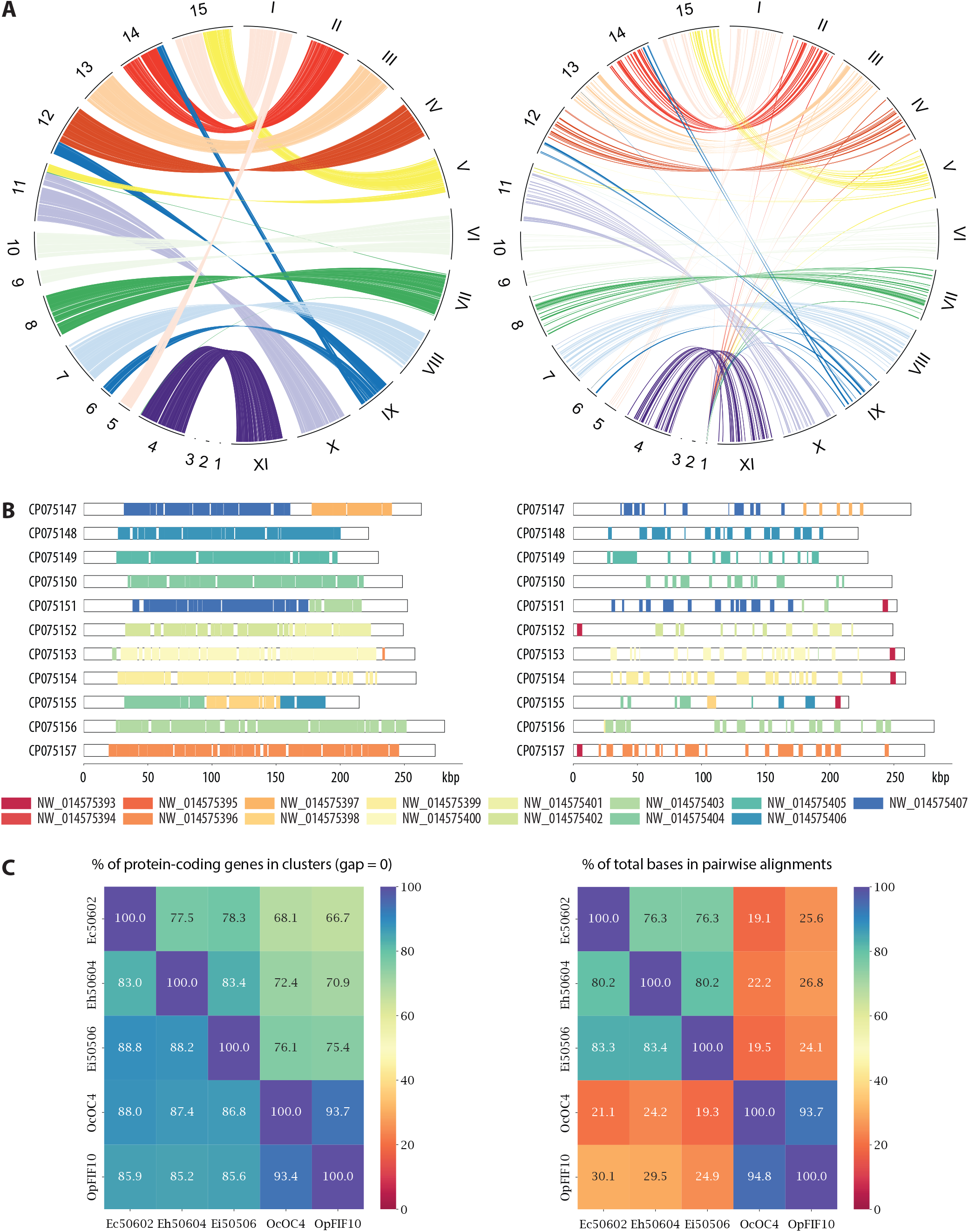
Examples of collinear maps generated with SYNY. Plots shown are from gene cluster (left) and genome alignment (right) inferences between the *Encephalitozoon hellem* ATCC50604 and *Ordospora colligata* OC4 genomes. (**a**). *Circos plots*. The reference contigs (from *E. hellem*) are labelled with roman numerals. The other contigs (from *O. colligata*) are labelled with Arabic numerals. Ribbons are color-coded based on the reference contigs. Plots were produced with the *label_size 80, no_ntbiases*, and *no_cticks* options. (**b**). *Chromosome maps (barplots)*. Contigs from *E. hellem* are labelled with their NCBI accession numbers (CP075147 to CP075157) and plotted to scale. Collinear regions with the *O. colligata* genome are highlighted by rectangular patches color-coded based on it contigs (NW_014575399 to NW_014575407). For brevity, only one of the identical legends between the left and right plots was kept. This legend was repositioned at the bottom of the panel to limit its width. (**c**). Heatmap summaries of proteins found in clusters (left) and of bases found aligned in collinear segments (right) between the *Encephalitozoon* and *Ordospora* genomes: Ec, *E. cuniculi*; Eh, *E. hellem*; Ei, *E. intestinalis*; Oc, *O. colligata*; Op, *O. pajunii*.

## Discussion

When we initially designed SYNY, we set out to develop a simple tool that identifies the gene clusters that are shared between eukaryote genomes and then plots this information as useful figures. To do this, we began by identifying gene pairs arrayed identically between genomes, then reconstructed gene clusters based on these gene pairs. We restricted our searches to protein coding genes to enable searches across larger evolutionary time scales but also to bypass RNA coding genes that are often unannotated in eukaryote genome projects. However, because the annotations used as inputs were sometimes missing protein-coding genes, featured multi-exon proteins annotated as separate entities, and/or contained spurious open reading frames annotated as hypothetical proteins, gene clusters inferred by SYNY were often fragmented due to these issues. To account for these possibilities, we implemented the option to allow for gaps between genes when performing cluster-based collinearity inferences. Likewise, to handle instances where genome annotations are not available, for example when comparing a newly assembled genome to one or more reference(s), we implemented a parallel approach based on genome sequences that leverages the Minimap2 [6] pairwise alignment tool.

One of our main goals with SYNY was to automate the production of collinearity plots that are both informative and suitable as is (or with as few modifications as possible) for publications purposes. To this end, we implemented some of the most common visualisation graphs in comparative genomics, including Circos [8] plots. Albeit SYNY runs relatively quickly on modest hardware, most of its runtime is dedicated to Circos plotting. Circos is not multithreaded, and while we parallelized its plotting instances across multiple threads (at the expense of using more random-access memory; RAM), generating the corresponding plots takes a substantial fraction of the total SYNY runtime. Because the number of pairwise comparisons to be performed can quickly balloon up with the number of genomes being compared, users may want to skip the Circos plots and use the barplots for preliminary visual investigations. The barplots are generated much faster (plotting 40 graphs from the case example took only 7 seconds for the barplots compared to 195 seconds for Circos) and run in a much smaller memory footprint.

The second most time-consuming segment of the SYNY pipeline involves the pairwise alignment step with Minimap2. Aligning larger genomes will lengthen is runtime and, because Minimap2 does not work with queries with larger than 2,147,483,647 bases (as per its user manual), inferring collinearity between genomes using pairwise alignments is not possible for queries exceeding that threshold. Aligning larger genomes with Minimap2 will also require more RAM. Performing a pairwise alignment between two 150 Mbp genomes from *Arabidopsis* peaked at about 10 Gb of RAM (0.7 Gb operating system (OS) + 9.3 Gb Minimap2; result not shown) whereas running gene cluster inferences on the same two genomes never exceeded 2.5 Gb of RAM. Considering the above and that cluster-based inferences perform better than pairwise alignments methods when dealing with high levels of sequence divergence (as shown in Fig. 2), users may want to skip pairwise alignments when dealing with well annotated genomes.

## Author contributions

JFP designed the pipeline; JFP and ATJ wrote the software and tested the pipeline; JFP and ATJ wrote and reviewed the manuscript.

## Funding

This work was supported in part by the National Institute of Allergy and Infectious Diseases of the National Institutes of Health [grant number R15AI128627] to JFP. The content is solely the responsibility of the authors and does not necessarily represent the official views of the National Institutes of Health.

## Notes

https://github.com/PombertLab/SYNY

### Competing Interest Statement

The authors have declared no competing interest.

